# Imprinting but not cytonuclear interactions affects parent-of-origin effect on seed size in *Arabidopsis* hybrids

**DOI:** 10.1101/2023.09.15.557997

**Authors:** Viviana June, Xiaoya Song, Z. Jeffrey Chen

## Abstract

The parent-of-origin effect on seed size can result from imprinting or a combinational effect between cytoplasmic and nuclear genomes, but their relative contributions remain unknown. To discern these confounding effects, we generated cytoplasmic-nuclear substitution (CNS) lines using recurrent backcrossing in the *Arabidopsis thaliana* ecotypes Col-0 and C24. These CNS lines differ only in the nuclear genome (imprinting) or in the cytoplasm. The CNS reciprocal hybrids with the same cytoplasm display a ∼20% seed size difference as observed in the conventional hybrids. However, seed size is similar between the reciprocal cybrids with fixed imprinting. Transcriptome analyses in the endosperm of CNS hybrids using laser-capture microdissection have identified 104 maternally expressed genes (MEGs) and 90 paternally-expressed genes (PEGs). These imprinted genes are involved in pectin catabolism and cell wall modification in the endosperm. *HDG9*, an epiallele and one of 11 cross-specific imprinted genes, controls seed size. In the embryo, a handful of imprinted genes is found in the CNS hybrids but only one is expressed higher in the embryo than endosperm. *AT4G13495* encodes a long-noncoding RNA (lncRNA), but no obvious seed phenotype is observed in the lncRNA knockout lines. *NRPD1*, encoding the largest subunit of RNA Pol IV, is involved in the biogenesis of small interfering RNAs. Seed size and embryo is larger in the cross using *nrpd1* as the maternal parent than in the reciprocal cross. In spite of limited ecotypes tested, these results suggest potential roles of imprinting and *NRPD1*-mediated small RNA pathway in seed size variation in hybrids.

## INTRODUCTION

Seed size is important to plant evolution and crop production and often show heterosis. Heterosis is a widely observed phenomenon in which the offspring show greater growth and fitness than either or both parents. Heterosis has been applied in agriculture to generate higher yielding and more resilient crops (Duvick, 2001; Lippman and Zamir, 2007; Chen, 2013; Schnable and Springer, 2013; Springer and Schmitz, 2017). The phenomenon was systematically studied by Charles Darwin, who observed that cross-pollinated plants displayed increased growth compared to self-pollinated plants (Darwin, 1876). And many studies and theories have attempted to explain the molecular basis that underlie this phenomenon (Lippman and Zamir, 2007; Chen, 2013; Schnable and Springer, 2013; Springer and Schmitz, 2017). An interesting observation is the parent-of-origin effect on heterosis where the offspring of the reciprocal crosses show varying degrees of heterosis (Waters et al., 2011; Miller et al., 2012; Groszmann et al., 2014; Ng et al., 2014). In animals, mules (horse x donkey, by convention, the maternal parent is listed first in a genetic cross) are larger and stronger than hinnies (donkey x horse) (McLean et al., 2019). The underlying mechanism for this is poorly understood. It involves imprinting (unequal expression of paternal and maternal alleles) in the nuclear genome and/or maternal effect from the cytoplasm. In *Arabidopsis* intraspecific hybrids, the parent-of-origin effect on biomass heterosis is mediated by the RNA-directed DNA methylation (RdDM) pathway that modulates levels of CHH (H=A, T, or C) methylation in the promoter of *Circadian Clock Associated1* (*CCA1*) (Ng et al., 2014), a central regulator of plant circadian clock (Greenham and McClung, 2015). This supports the notion that imprinting mediates the parent-of-origin effect on hybrid vigor (Vu et al., 2013; Ng et al., 2014). However, in the conventional reciprocal crosses, maternal effect cannot be ruled out simply because the effects of imprinted gene expression in the nuclear genome and cytoplasmic genomes are confounded in the reciprocal hybrids.

The cytoplasm consists of mitochondrial and plastid genomes that affect plant growth and development. In wheat alloplasmic lines, cytoplasmic genomes from *Aegilops* affects flowering time and fertility in wheat (Kihara, 1982). Additional studies of these lines have found that substitution of the cytoplasm is associated with a variety of changes in the expression of genes involved in photosynthetic components, mitochondrial electron transport chain, and retrograde signaling to the chloroplast (Crosatti et al., 2013). In maize, substitution of maize cytoplasm with cytoplasm from the wild progenitor *teosinte* leads to a decrease in yield, and cytonuclear interactions can change plant height in F_2_ populations (Edwards et al., 1996; Tang et al., 2013). In *Arabidopsis*, cytonuclear interactions contribute to changes in the metabolome and growth (Douglas et al., 2015; Joseph et al., 2015). In cybrids generated by haploid inducer line in six genotypes (Bur, C24, Col-0, L*er*, Sha, WS-4, and ELy), certain plasmotypes, particularly that of the accessions Bur and Ely, show most epistatic effects (Flood et al., 2020). Many of these cybrids have a strong effect on photosynthetic phenotypes (Joseph et al., 2015; Flood et al., 2020), as cytoplasmic genomes play an important role in photosynthesis and growth.

Imprinted genes in the nuclear genome may also play a role in the parent-of-origin effect (Botet and Keurentjes, 2020). According to the parental conflict theory, maternally expressed imprinted genes may limit the growth of offspring and conserve more maternal resources, whereas paternally expressed imprinted genes tend to drive growth of the offspring and promote the allocation of maternal resources towards this growth (Haig, 2000, 2013). In plants, imprinted genes are known to be involved in regulating endosperm proliferation and seed growth. Studies in maize, rice, and *Arabidopsis* have identified a cross-specific imprinting phenomenon (Waters et al., 2013; Pignatta et al., 2014). For example, *HDG3* is a paternally expressed imprinted gene in the majority of *Arabidopsis* accessions, but not in Cvi (Pignatta et al., 2014). These imprinted genes in reciprocal hybrids likely contributes to seed development.

Many imprinted genes and their roles in seed development have been studied in *Arabidopsis* (Gehring et al., 2011; Hsieh et al., 2011; Raissig et al., 2013; Pignatta et al., 2014; Fort et al., 2017). However, the contribution of cytoplasmic genomes and imprinted genes in the nuclear genomes to the seed development has not been separated in these studies.

To discern these confounding effects, we generated cytoplasmic-nuclear substitution (CNS) lines. The nuclear genome of the maternal parent is replaced by 6 generations of backcrossing with the recurrent (paternal) parent, while the cytoplasm remains the same. These CNS lines are also known as alloplasmic lines in wheat (Kihara, 1951) or *Brassica* (Bonhomme et al., 1992), and are often used to generate cytoplasmic-nuclear male sterility lines in plants, due to the incompatibility between cytoplasmic and nuclear genomes. Here, we created a pair of CNS lines using the *Arabidopsis thaliana* Col and C24 ecotypes through six generations of backcrossing, and the nuclear genome was 99.99% identical to that in the maternal parent. The seedling phenotypes of CNS lines resembled the paternal parent. Interestingly, the parent-of-origin effect on seed size was found only in the reciprocal CNS crosses where the cytoplasm is the same. On the contrary, in the reciprocal crosses with the same nuclear genome combination, the parent-of-origin effect disappeared. We further characterized the imprinted gene expression using laser-capture microdissection (LCM) in the reciprocal crosses with the same cytoplasm. Despite one pair of genotypes is tested, the results indicate a role of imprinted genes of the nuclear genome in the seed size heterosis in the CNS reciprocal hybrids. Moreover, the seed size difference is also associated with the small RNA pathway involving *NRPD1*.

## RESULTS

### Generation and genotype conformation of CNS lines through a backcrossing scheme

Using a backcross scheme (Figure 1A), we obtained the CNS lines Col(C24C) by backcrossing the *A. thaliana* accessions Col with pollen from C24C for six generations (expected homozygosity of 99.98%) and the CNS lines C24(ColC) by backcrossing C24 with pollen from ColC for five generations (∼99.95% of homozygosity). We genotyped multiple plants of each CNS line using simple sequence repeats and selected four lines (two of each CNS line), two parental lines, and one F_1_ hybrid (C24 x Col) for sequencing to determine percentage of homozygosity (Figure 1B, Figure S1, A and B). Perfectly mapped sequence reads onto TAIR10 genome were used to call variants from other genotypes using GATK’s HaplotypeCaller (Van der Auwera et al., 2013). A total of 591,232 SNPs were unique to either the Col or the C24 genome and present as heterozygous in the F_1_ (C24 x Col). We found that >99.9% of the SNPs were homozygous for the genome in the Col(C24C) and C24(ColC) CNS lines, respectively (Figure 1B and Figures S1, A and B), as expected after five to six generations of backcrossing.

**Figure 1.**
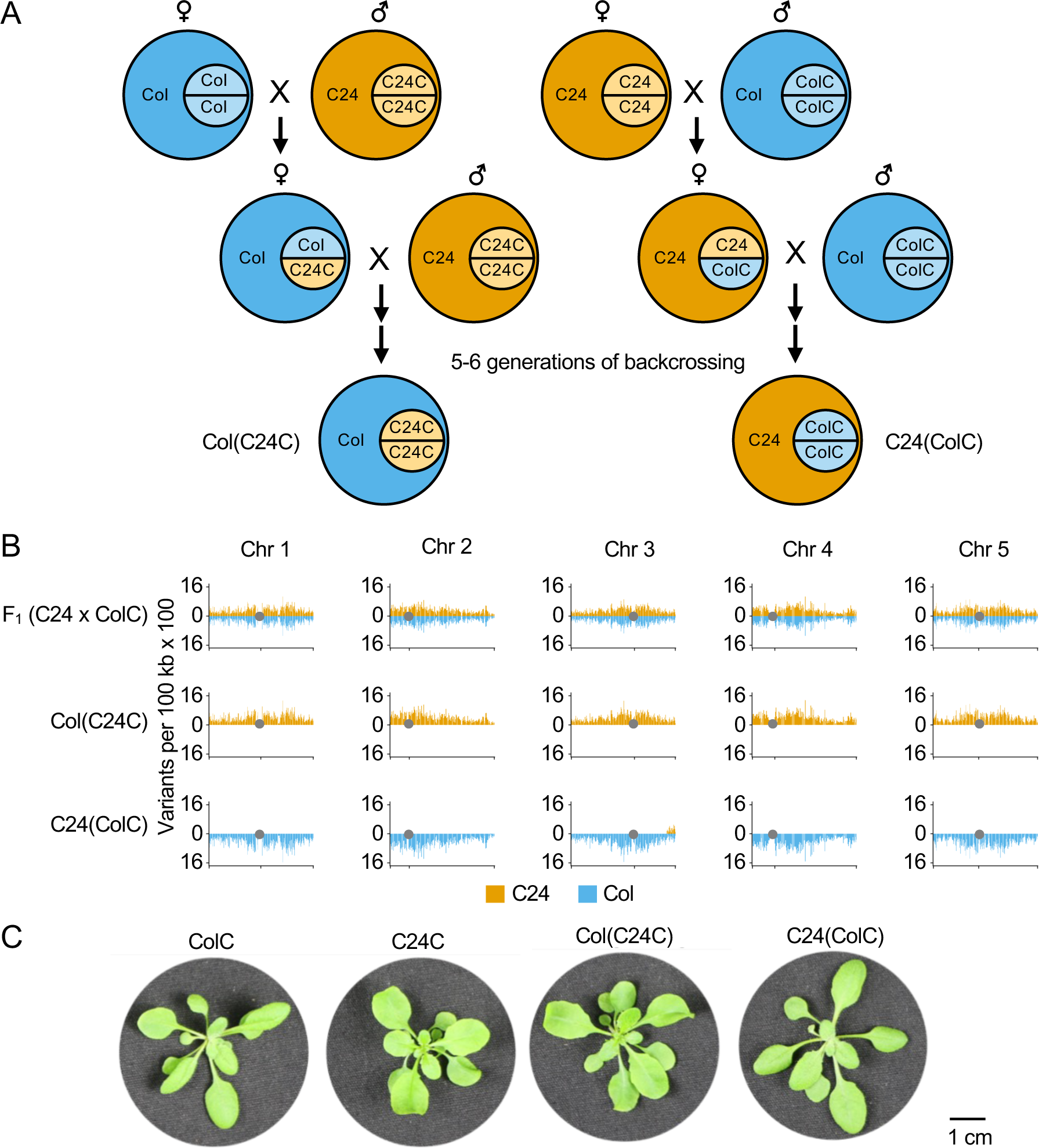
Generation of cytoplasmic-nuclear substitution (CNS) lines and genomic analysis. (A) Backcrossing scheme used to generate the CNS lines, Col(C24C) and C24(ColC) lines, in which the nucleus in respective parental lines is replaced by C24C and ColC, respectively. C refers to the *CCA1:LUC* reporter in the C24 or Col line. (B) SNPs called in the CNS lines that are unique to either the C24 or Col genome, confirming the conversion of the nuclear genome. Heterozygous F1(C24 x ColC) is shown as a control. SNPs are grouped into 100 kb bins. Grey circles indicate the position of the centromere. (C) Typical rosette images from the parental and CNS lines at 21 days after sowing. Scale bar = 1 cm for all images.

The chloroplast and mitochondrial genomes were analyzed by PCR using known polymorphic markers to confirm retention of the cytoplasmic genomes. PCR was used to amplify a polymorphic region upstream of the *atp6-2* gene in the mitochondrial genome (Forner et al., 2008) and a polymorphic region between *rbcL* and *accD* in the chloroplast genome (Azhagiri and Maliga, 2007). Using these polymorphic markers, both the Col(C24C) and C24(ColC) lines were confirmed to have retained their original maternal mitochondrial and plastid genomes, respectively (Figure S1, C and D). Taken together with the nuclear genome sequencing results, the Col(C24C) and C24(ColC) CNS lines were converted as expected for further analyses.

### CNS lines resembled the lines with the same nuclear genome, and the seed size variation depends on the nuclear genome

Using the CNS lines, we investigated changes in phenotypic traits that are caused by different combination of nuclear and cytoplasmic genomes. We found that seedling phenotypes resembled that of their paternal parent. For example, Col(C24C) resembled the C24 accession, and C24(ColC) resembled the Col-0 parent (Figure 1E). There was no difference in flowering time (Figure S1E), rosette diameter (Figure S2A), or seed size (Figure S2B and Figure S2C) between the CNS lines and their paternal parents. This is in agreement with the notion that most cytoplasmic swaps did not change these phenotypes (Flood et al., 2020).

In previous studies C24xCol hybrids displayed increased biomass vigor as compared to the ColxC24 hybrids (Groszmann et al., 2014; Ng et al., 2014; Miller et al., 2015). There was 20-30% difference in mature seed size between the Col x C24 and C24 x Col hybrids (Miller et al., 2012; Groszmann et al., 2014). However, these studies cannot discern the effect of cytoplasmic and nuclear genomes on the seed size. Using the CNS lines, we generated pairs of reciprocal hybrids that have either the same cytoplasmic genomes or the same nuclear genome to separate these effects in the reciprocal hybrids (Figure 2A). In the reciprocal CNS hybrids Col(C24C) x Col and Col x C24C, which have the same cytoplasmic genomes, seed size was ∼20% larger in the Col(C24C) x Col cross than in the reciprocal cross Col x Col(C24C); this resembled the seed size difference between the conventional reciprocal hybrids (C24C x Col and Col x C24C) (Figure 2B, Figure S2B). On the contrary, no difference in the mature seed size was observed between the reciprocal cybrid crosses Col(C24C) x Col and C24C x Col with the same nuclear genome combination but different cytoplasmic genomes.

**Figure 2.**
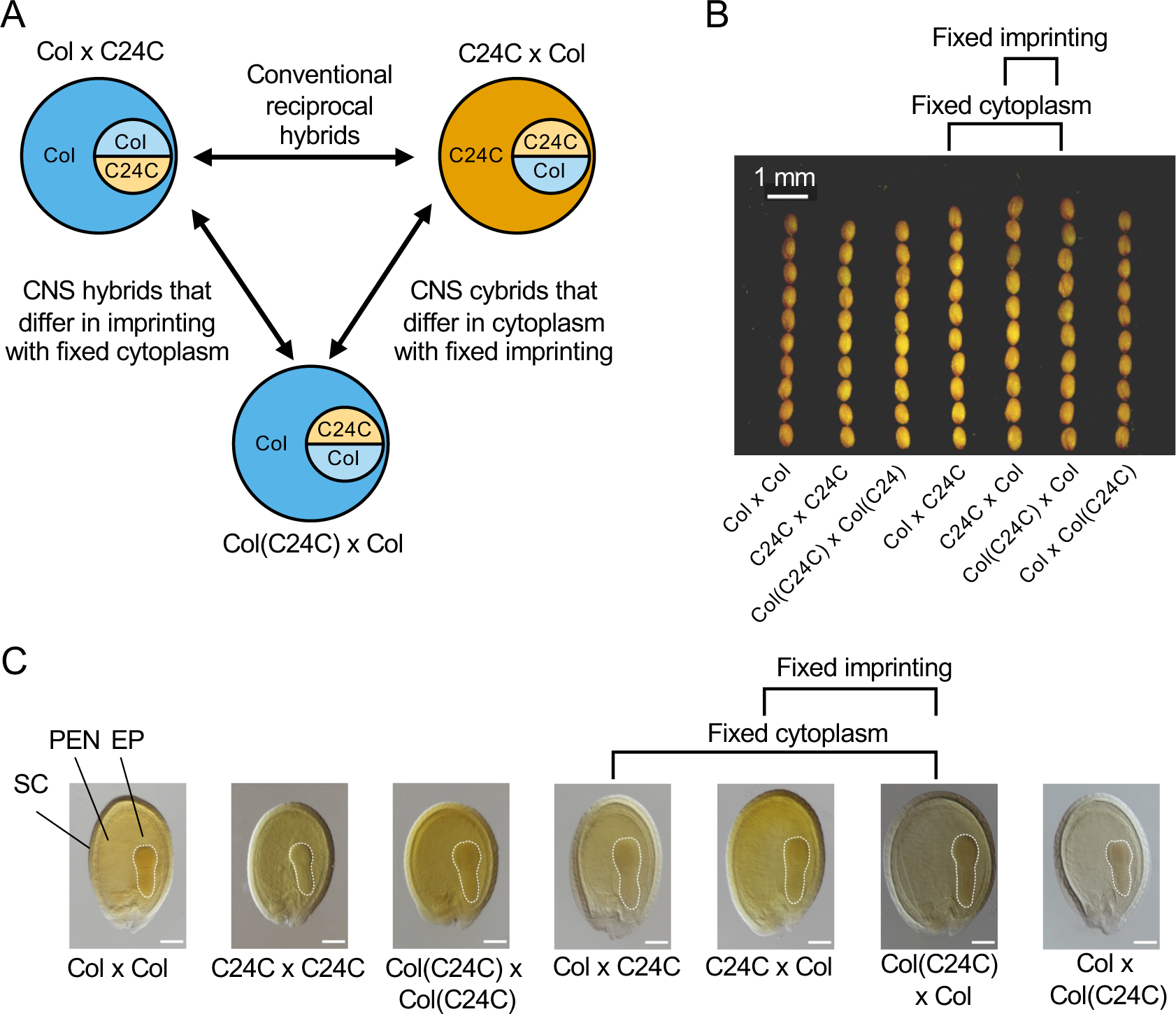
Parent-of-origin effects on seed size in CNS and conventional reciprocal hybrids. (A) The genotypic differences between conventional hybrids, CNS hybrids with fixed cytoplasm, and CNS cybrids with fixed imprinting. (B) Mature seeds are larger in the C24C x Col than in the Col x C24C cross. This difference was observed in CNS reciprocal hybrids with fixed cytoplasm, but not in the CNS reciprocal cybrids with fixed imprinting. Scale bar = 1 mm for all images. (C) Micrographs of cleared seeds at 7 days after pollination (DAP), showing the embryo proper (EP), peripheral endosperm (PEN), and seed coat (SC). Scale bar = 100 µm for all images.

This seed size difference was established early during seed development. At seven days after pollination (DAP), the seed size was ∼20% larger in Col(C24C)xCol and in C24xCol than in the ColxCol(C24C) or ColXC24 crosses (Figure 2C, Figure S2C). The seed size was similar between the reciprocal cybrids where nuclear genome combination was the same. This result indicates that imprinting alone determines seed size and endosperm development.

### Imprinted genes identified in the endosperm between CNS reciprocal hybrids

To investigate how imprinted gene expression changes during seed development, we isolated embryos and endosperm from seeds at 6 DAP using laser capture microdissection (LCM) (Belmonte et al., 2013; Ando et al., 2021). These LCM samples were used for mRNA-seq analysis to identify imprinted genes using a generalized linear model (Wyder et al., 2019).

Using 214,274 informative SNPs from the Col(C24C) line relative to the Col reference, we evaluated allelic expression of 15,134 loci in the endosperm. Our dataset was also free of seedcoat RNA contamination (Figure S3C and Figure S3D), a problem present in several previous studies of imprinted genes (Schon and Nodine, 2017). In the conventional reciprocal crosses, we identified 123 maternally expressed genes (MEGs), and 108 paternally expressed genes (PEGs) (Figure 3A) (Supplementary Dataset 2). In reciprocal crosses with the same cytoplasm, we found 104 MEGs and 90 PEGs (Figure 3B) (Supplementary Dataset 2). The overlap between the two sets of data were very high, and 99 MEGs and 89 PEGs, respectively, were shared in both conventional and fixed-cytoplasm reciprocal hybrids (Figure 3, C and D). However, very few imprinted genes were identified in the reciprocal cybrids but not in the conventional reciprocal crosses. These data indicate that the cytoplasmic genomes play a minor role in imprinted gene expression. Moreover, the imprinted genes that were specific to the conventional or cybrid reciprocal crosses tend to show weaker expression levels than MEGs and PEGs shared in both sets of crosses (Figure S4).

**Figure 3.**
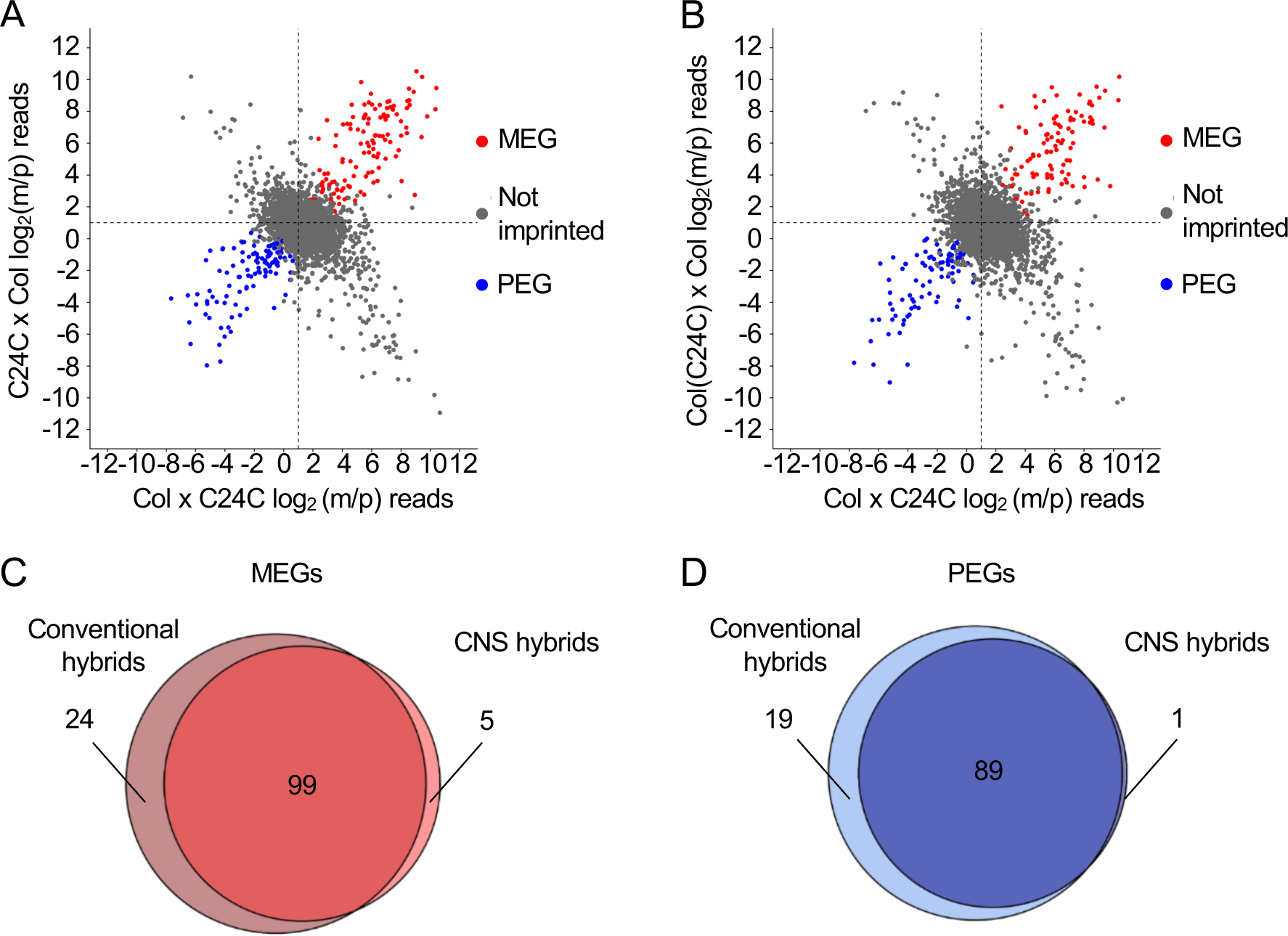
Comparative analysis of imprinted genes in the CNS and conventional reciprocal hybrids. (A) Expression plots of maternal and paternal reads (mean of 2 biological replicates) from the conventional reciprocal crosses (C24C x Col, *y*-axis, and Col x C24C, *x*-axis). Red, blue, and grey spots indicate maternally-expressed genes (MEGs), paternally-expressed genes (PEGs), and not imprinted genes, respectively. (B) Expression plots of maternal and paternal reads (mean of 2 biological replicates) from the CNS reciprocal crosses with fixed cytoplasm [Col(C24C) x Col, *y*-axis, and Col x C24C, *x*-axis]. Red, blue, and grey spots indicate MEGs, PEGs, and not imprinted expression, respectively. (C, D) Overlap of MEGs (C) or PEGs (D) in the conventional hybrids and CNS hybrids. Source data: Dataset S2, endosperm imprinting.

Gene Ontology (GO) analysis found that the genes upregulated in the Col x C24C cross relative to the Col(C24C) x Col cross were enriched for cell wall modification (GO:0042545) and carbohydrate metabolic process (GO:0005975) (Figure S7B). They included many pectin methylesterase genes, which were found to be upregulated in the linear cotyledon stage of seed development (Wolff et al., 2015). It is likely that the Col x C24C seeds are developing slightly faster than the reciprocal cross, resulting in the increased expression of these genes that typically are upregulated around 6 DAP. The delay of entering this phase in the reciprocal cross Col(C24C) x Col may prolong cellularization, allowing the seeds to grow larger.

Our study identified approximately half of the PEGs and two-thirds of the MEGs, respectively (Figure 4), as previously reported (Gehring et al., 2011; Hsieh et al., 2011; Pignatta et al., 2014). This degree of variation is not unexpected, as there is a considerable amount of variation between imprinted genes identified among different studies (Klosinska et al., 2016; Schon and Nodine, 2017; Wyder et al., 2019). This may be related to CNS lines and different ecotypes used, contamination of the endosperm with seed coat (Schon and Nodine, 2017), and/or different statistical methods and cut-off values used for data analysis (Wyder et al., 2019).

**Figure 4.**
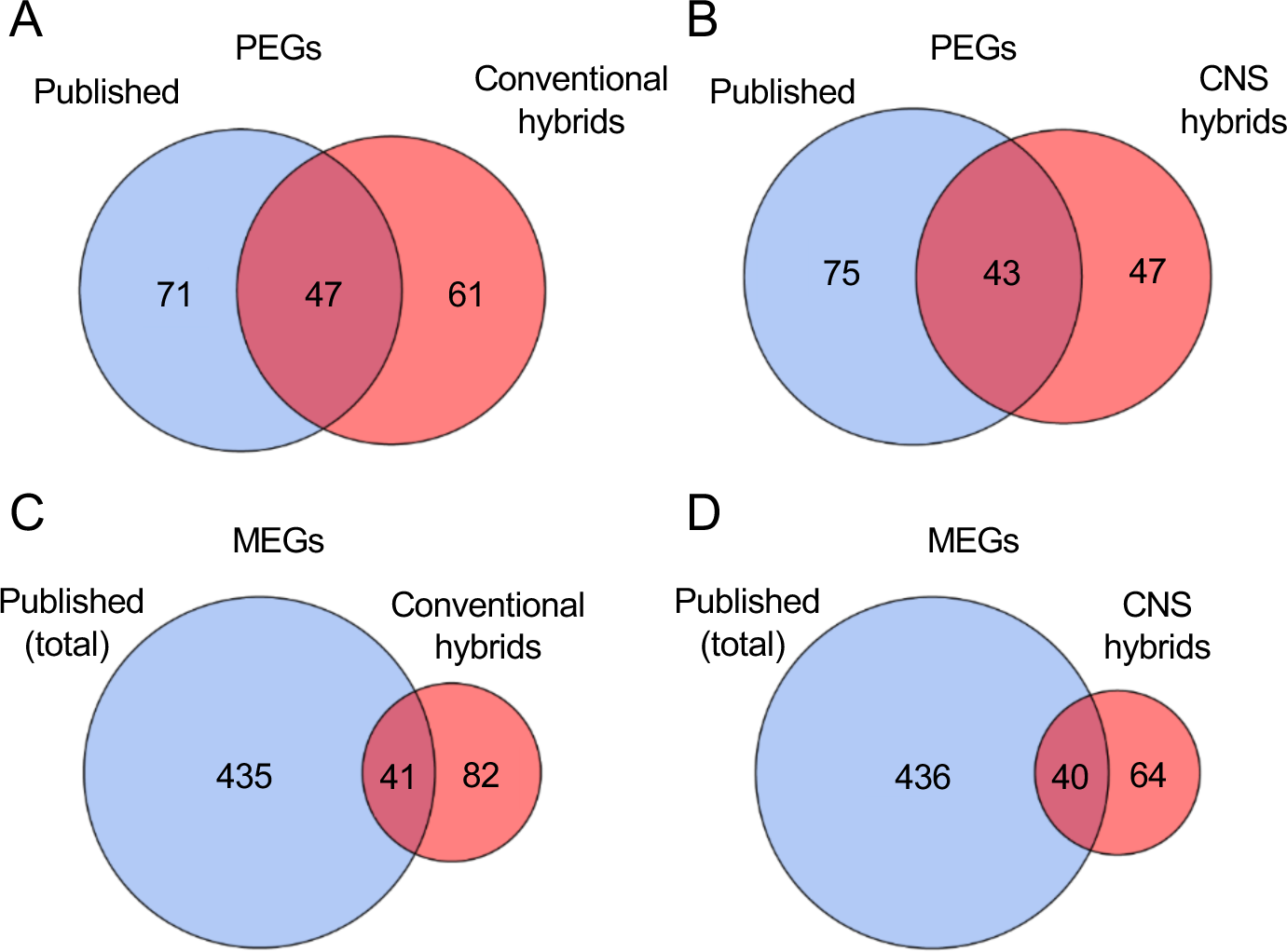
Comparative analysis of the imprinted genes identified from this and other studies. (A, B) Overlap between PEGs identified in the published studies (Hsieh 2011, Gehring 2011 and Pignatta 2014) and the PEGs identified in this study in the conventional reciprocal hybrids (A) and CNS reciprocal hybrids (B). (C, D) Overlap between the MEGs identified in the published studies (Hsieh 2011, Gehring 2011 and Pignatta 2014) and those identified in this study in the conventional reciprocal hybrids (C) and CNS reciprocal hybrids (D).

### Cross-specific imprinting in the endosperm between the CNS reciprocal hybrids

Epigenetic variation contributes to changes in gene expression across *Arabidopsis* accessions (Kawakatsu et al., 2016), as the presence of epialleles in certain accessions can eliminate or establish imprinting during seed development (Hsieh et al., 2011; Pignatta et al., 2014). Seed size variation between *Arabidopsis* accessions could be related to cross-specific imprinting in reciprocal hybrids (Pignatta et al., 2014; Pignatta et al., 2018). Cross-specific imprinted genes lack imprinting in one of the two reciprocal crosses between some ecotypes.

Here, we identified eleven genes that showed biased expression in one cross, but not in the reciprocal cross, indicating an epiallele present in only one parental accession (Figure 5 and Supplementary figure S3). Three MEGs lacked maternal bias in the crosses using C24 as the maternal parent (Figure 5B), three PEGs lost paternal bias in the crosses using C24 as the paternal parent (Figure 5C), and four MEGs lacked maternal bias in the crosses using Col as the maternal parent (Figure 5D) (Supplementary Dataset 4).

**Figure 5.**
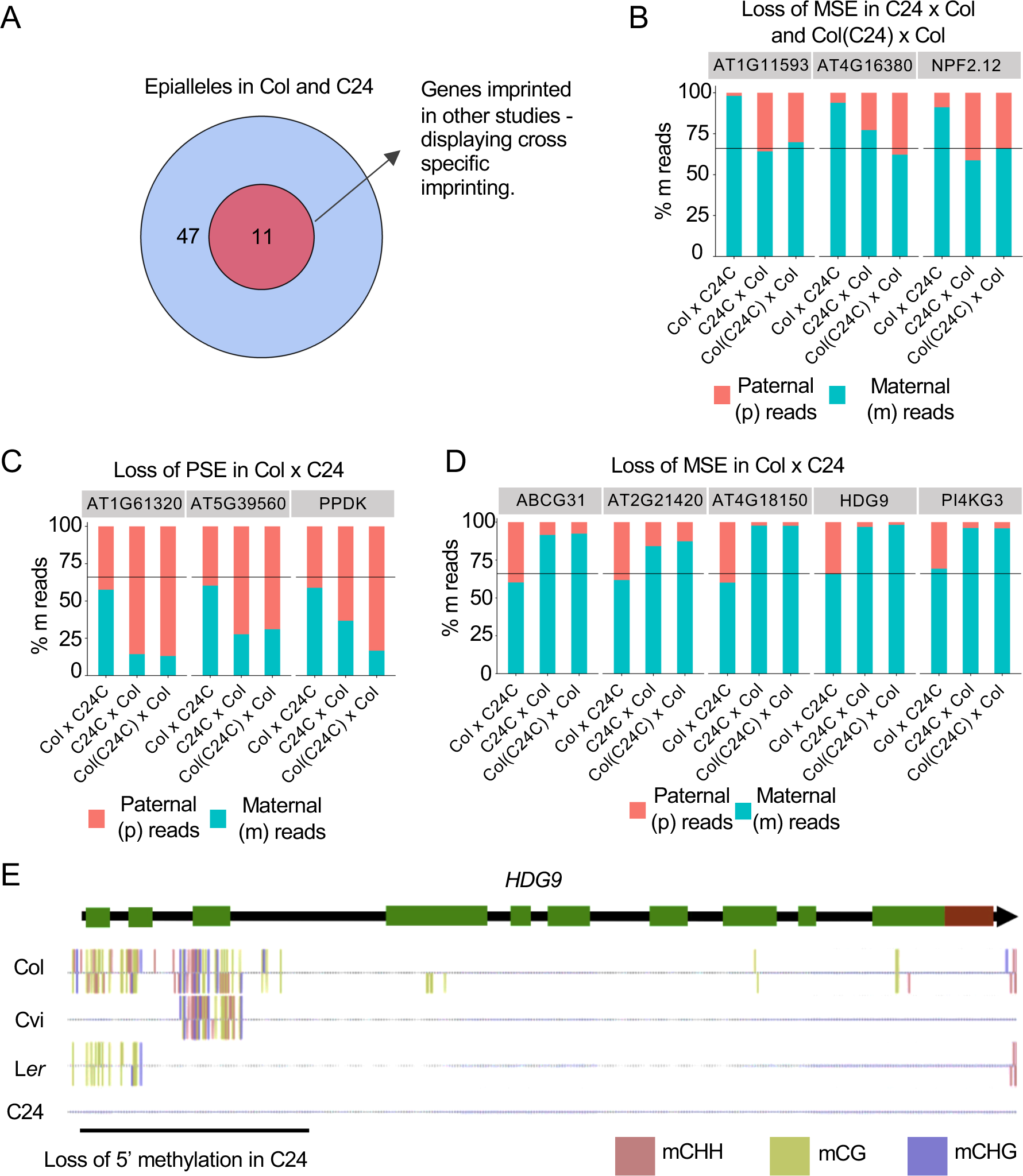
Analysis of cross-specific imprinted genes in the endosperm. (A) There were 58 epialleles in Col and C24, 11 of which were identified as imprinted genes in previous studies (Hsieh et al. 2011, Gehring et al. 2011 and Pignatta et al. 2014). (B) Three imprinted genes displaying loss of maternal-specific expression (MSE) in C24 x Col and Col(C24) x Col. (C) Three imprinted genes displaying loss of paternal-specific expression (PSE) in Col x C24. (D) Five imprinted genes displaying loss of MSE in Col x C24. (E) Methylation patterns on *HDG9* in Col, CVI, L*er* and C24 from the 1,001 Epigenomes Project. Source data: Datasets S3, cross specific imprinting counts.

The epiallelic variation in cross-specific imprinting could be related to DNA methylation. We examined eleven genes using the data from the 1,001 epigenomes project (Kawakatsu et al., 2016). Surprisingly, only one gene *HDG9*, encoding a homeodomain glabrous (HDG) transcription factor, showed allelic methylation patterns (Figure 5E). The 5’ region of *HDG9* was differentially methylated, and this methylation was lost in C24. In the crosses where *HDG9* is imprinted, the maternal allele is demethylated, while the paternal allele remains methylated. This change in methylation is coincident with the loss of imprinting observed in the Col x C24C hybrid. The 5’ methylation could silence the paternal allele, and loss of this methylation in C24 correlated with increased *HDG9* expression levels in the cross using C24 as the paternal parent.

HDG transcription factors are involved in endosperm development (Pignatta et al., 2018) and act antagonistically with AINTEGUMENTA-LIKE (AIL) transcription factors to regulate cell proliferation in several plant tissues (Horstman et al., 2015). AILs generally promote cell proliferation, while HDGs restrict it. Thus, a change in imprinting in *HDG9* could contribute to the seed size differences that were observed between the reciprocal hybrids involving C24. *HDG3* is a cross-specific imprinted gene in reciprocal crosses involving the Cvi accession (Pignatta et al., 2018), and it is possible that a similar mechanism underlying *HDG9* contributes to the seed size difference in the Col x C24 reciprocal crosses. To test this, we generated *HDG9* knockouts in a C24 and Col-0 background using CRISPR/Cas9 and a cassette of the two gRNAs (Xing et al., 2014) and the pHEE401E construct (Wang et al., 2015). These lines were used to test the effect of *HDG9* on seed size (Figure S6, A and B). Knockout of *HDG9* in C24, but not in Col-0 led to an increase in mature seed size, suggesting the effect of the *HDG9* knockout is limited to the cross-specific epiallele in the C24 ecotype.

### Cytoplasmic genomes contribute little to the seed size heterosis

Seed size was similar in the reciprocal crosses where the nuclear genome combination (imprinting) is the same or fixed, suggesting little contribution of the cytoplasmic genomes to the parent-of-origin effect on seed size heterosis. Indeed, only ten differentially expressed genes in the endosperm were identified when the cytoplasmic genomes were fixed (Figure S7 and Supplementary Dataset 4). Moreover, these differentially expressed genes belonged to those coded by the cytoplasmic genome itself, such as cytochrome oxidase subunit C (*COX3*), NAD(P)H quinone oxidoreductase K and NAD(P)H quinone oxidoreductase C, which were upregulated in the lines or crosses carrying a C24 cytoplasm. This is consistent with allelic variation between Col and C24, as reported (Forner et al., 2005), and the *cox3* locus generates additional *COX3* transcript isoforms associated with the C24 cytoplasm.

### Imprinted genes identified in the embryo between the CNS reciprocal hybrids

In contrast to the endosperm, relatively few genes were imprinted in the embryo. In the embryo of conventional reciprocal crosses, we identified seven MEGs and three PEGs (Figure 6A and Supplementary Dataset 6). In the embryo of the reciprocal crosses with the same cytoplasm, we identified four MEGs and one PEG (Figure 6B and Supplementary Dataset 6). Again, these imprinted genes in the embryo overlapped between reciprocal crosses of the conventional lines and CNS lines (Figure 6C). Notably, all MEGs and one of the PEGs identified in the embryo were expressed at higher levels in the endosperm than in the embryo (Figure S6). This could be due to a potential contamination from the endosperm tissues in the embryo (Schon and Nodine, 2017). However, our LCM approach has eliminated the contamination in the embryo tissues (Figure S4).

**Figure 6.**
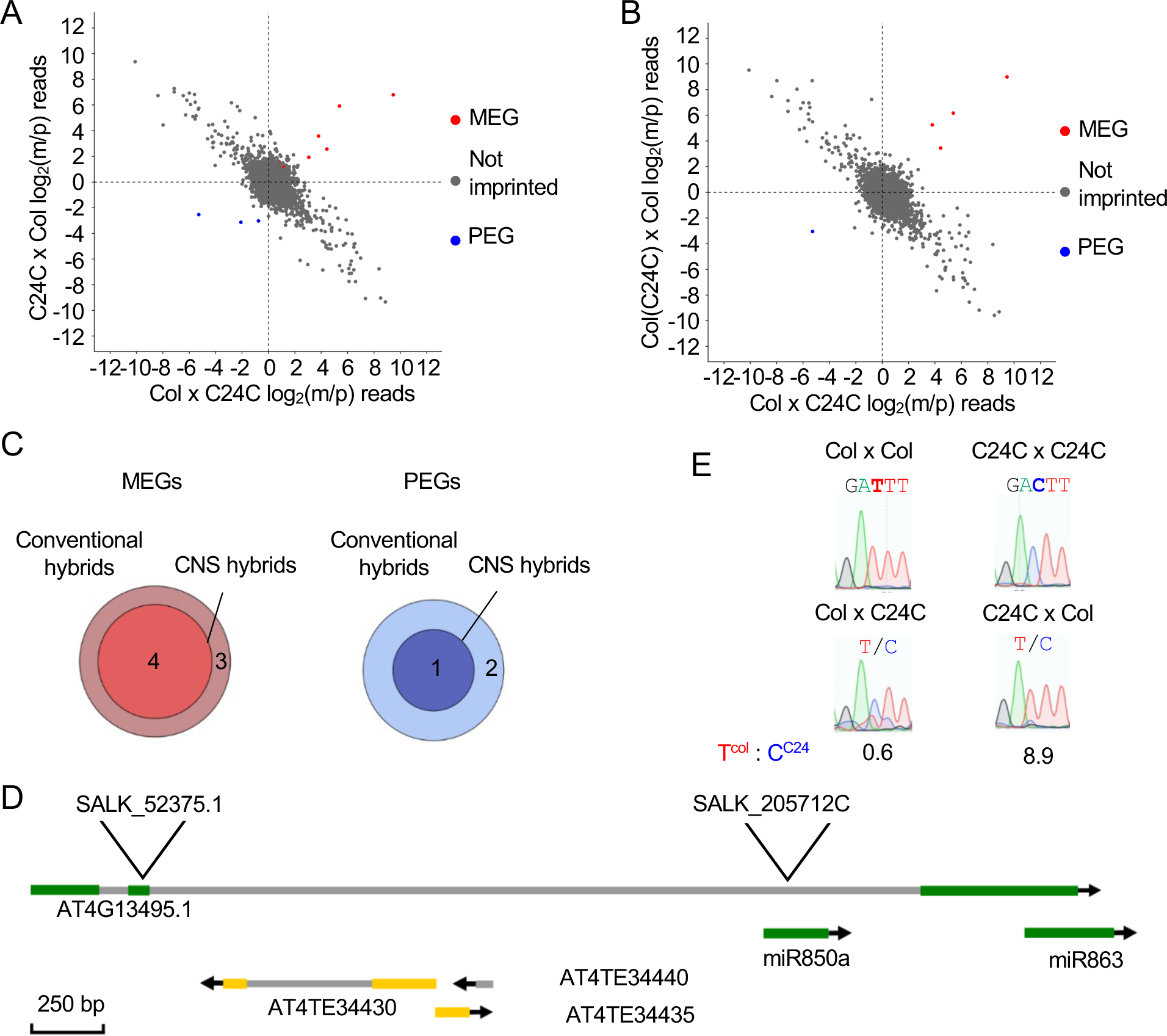
Analysis of the paternally expressed gene AT4G13495 in the embryo. (A) Expression plots of maternal and paternal reads (mean of 2 biological replicates) from the conventional reciprocal crosses (C24C x Col, *y*-axis, and Col x C24C, *x*-axis). Red, blue, and grey spots indicate maternally-expressed genes (MEGs), paternally-expressed genes (PEGs), and not imprinted genes, respectively. (B) Expression plots of maternal and paternal reads (mean of 2 biological replicates) from the fixed cytoplasm reciprocal crosses [Col(C24C) x Col, *y*-axis, and Col x C24C, *x*-axis]. Red, blue, and grey spots indicate MEGs, PEGs, and not imprinted expression, respectively. (C) Overlap of the imprinted genes between CNS hybrids and conventional hybrids. (D) Genomic feature of the locus AT4G13495. The lncRNA overlaps with two miRNAs, miR850 and miR863a. Source data: Dataset S3, embryo imprinting. (E) Expression validation of imprinted lncRNA AT4G13495 with Sanger sequencing. Ratios indicate the mean ratio of the Col allele to the C24 allele (*n* = 3). Source data: Dataset S6, embryo imprinting.

Only one PEG (*AT4G13945*) in the embryo displayed higher expression levels in the embryo than in the endosperm (Figure S7). *AT4G13945* encodes a long non-coding RNA (lncRNA) and has been identified as imprinted gene in the embryo of triploid Col x C24 crosses (Fort et al., 2017) and in the endosperm of reciprocal Col x L*er* hybrids (Gehring et al., 2011). Our data suggest that *AT4G13945* expression is embryo specific, with an expression level that is 10.9-fold higher in the embryo than in the endosperm at 6 DAP (Figure S8). It is likely that the presence of three transposable elements that lie within this locus inadvertently led to the imprinting of the full transcript in the embryo (Figure 6C).

*AT4G13495* lncRNA overlapped with two miRNA loci, miR863 and miR850 (Figure 6D). Both *AT4G13495* and miR863 showed paternally-biased expression in the embryo (Figure 6E). The lncRNA has been found to be involved in retrograde signaling to the chloroplasts (Habermann et al., 2020). In the vegetative tissues, *AT4G13495* transcripts from Col x C24 and C24 x Col were biallelically expressed in rosettes (Figure S9E), using a previously published RNA-seq dataset (Miller et al., 2015).

To test the effect of this lnRNA in seed size, we examined SALK T-DNA knockout lines. In both T-DNA insertion lines, no transcript was detected by RT-qPCR (Figure S9, B and C). However, no obvious phenotype was observed during embryogenesis or vegetative growth in these two lines (Figure S9, F and G). This lncRNA can moderate *SERRATE* expression through miR863 (Niu et al., 2016; Habermann et al., 2020). We evaluated the expression of miR863 and *SERRATE* in the mutants with RT-qPCR. Although miR863 was not expressed in the *at4g13495* mutants, *SERRATE* expression remained unchanged in the mutants (Figure S9D). The exact function of this lncRNA during embryo and seed development needs further investigation.

### Seed and embryo size heterosis is affected by *NRPD1*

*NRPD1* encodes the largest subunit of plant-specific RNA Pol IV and a homolog of the RNA polymerase II subunit and is involved in biogenesis of small interfering RNAs (siRNAs) (Herr et al., 2005; Onodera et al., 2005). Moreover, a group of *NRPD1*-dependent or p4-siRNAs is maternally transmitted (Mosher et al., 2009). Biogenesis of these p4-siRNAs is dependent on ploidy and maternal genome dosage in the endosperm (Lu et al., 2012), and they regulate expression of the target including some imprinted genes in the endosperm (Lu et al., 2012; Kirkbride et al., 2019). To test the effect of *NRPD1* on seed size, we compared mature seed size in the reciprocal crosses involving *nrpd1* (Figure S10, A and B). The seed size was substantially reduced in the *nrpd1* x C24 hybrid, compared to the reciprocal hybrid (C24 x *nrpd1*) or wild-type hybrid (C24 x Col). At 6 days after pollination (DAP), the embryo size was also larger in the *nrpd1* xC24 hybrid relative to the reciprocal hybrid (C24 x *nrpd1*) or C24 x Col. Together, these results indicate a potential role for *NRPD1* is in seed and embryo size.

## DISCUSSION

Maternal and paternal genomes are critical to the growth and development of offspring in sexually reproductive organisms like humans and flowering plants. However, the parent-of-origin effects in plants are confounded by the imprinting (nuclear genes) and cytonuclear genome interactions. Using the CNS lines, we can discriminate between the imprinting and cytoplasmic effects. Notably, cybrids can also be produced in various ecotypes (Flood et al., 2020) using haploid inducer techniques (Ravi and Chan, 2010). The latter may introduce some side effects by the centromeric protein and transgenic approach, while CNS lines may retain a minute amount of residual heterozygosity. Nonetheless, through generating CNS reciprocal hybrids that had either fixed cytoplasmic genomes or fixed imprinting, we found that the parent-of-origin effect on seed size is largely dependent on imprinting, while cytoplasmic genomes play a minor role in gene expression and seed size variation.

The MEGs and PEGs identified in the CNS reciprocal crosses overlap with many imprinted genes that were identified in previous studies (Gehring et al., 2011; Hsieh et al., 2011; Pignatta et al., 2014), confirming the role of imprinting in seed size variation. The cross-specific imprinted genes also contribute to the parent-of-origin effect (Pignatta et al., 2014; Pignatta et al., 2018). For example, in Cvi, a loss of methylation at *HDG3* removes imprinting of the gene resulting in biallelic expression, contributing to the small seed size of seeds with Cvi as the paternal parent (Pignatta et al., 2014). Similarly, *HDG9* is biallelically expressed when C24 is used as the paternal parent, but is a MEG in the crosses involving L*er*, Col and Cvi accessions. This lack of *HDG9* imprinting in C24 is like the lack of imprinting at *HDG3* in Cvi when Cvi is used as a parental parent (Pignatta et al., 2014; Pignatta et al., 2018). Both *HDG9* and *HDG3* are class IV homeodomain glabrous transcription factors that are likely involved in endosperm development. *HDG3* is thought to regulate endosperm cellularization; a decrease in *HDG3* expression leads to premature endosperm cellularization, limiting seed size and contributing to smaller seeds. We found that C24 is associated with larger seed size when it is used as the maternal parent and smaller seed size when it is used as the paternal parent. Knockout of *HDG9* in C24 led to an increase in mature seed size, suggesting the effect of the *HDG9* knockout is limited to the cross-specific epiallele in the C24 ecotype.

Using CNS lines in the reciprocal crosses, we found minor effects of the cytoplasmic genomes on seed size heterosis. A small set of genes that are affected by parent-of-origin effect are related to photosynthesis, such as NPQ and Φ_PSII_, as previously reported (Flood et al., 2020). This is probably because imprinting is limited to the seed and endosperm development in plants and the parent-of-origin effect is not common in plant somatic tissues. In addition, the genetic diversity between cytoplasmic genomes may affect cytoplasmic-nuclear compatibility, leading to cytoplasmic-nuclear male sterility (CMS). For example, ecotypes such as Ely with a mutation in *PsbA* that affects PSII efficiency (El-Lithy et al., 2005) show greater phenotypic effects among the cytoplasmic swap lines (Flood et al., 2020), including changes in biomass, flowering time, seed size, photosynthetic efficiency, and CMS. Among the hybrids derived from the ecotypes with a low level of cytoplasmic diversity, nuclear gene and genomic interactions determine imprinting and seed size phenotypes.

The imprinted genes identified in study are largely void of seedcoat contamination, a predominant problem as previously noted (Schon and Nodine, 2017). Using CNS lines and LCM, we also found a few imprinted genes in the embryo. Many of the imprinted genes in the embryo are also expressed at higher levels in the endosperm with one exception. The gene *AT4G13495* is strongly expressed in the embryo but weakly expressed in the endosperm. A previous investigation suggests that this lncRNA may be involved in retrograde signaling to the chloroplast (Habermann et al., 2020). Imprinting of this lncRNA is related to three adjacent TEs, while two miRNAs within the locus may control the expression of target genes. The role of this lncRNA in embryo and see development remains to be tested. Finally, the parent-of-origin effect on seed and embryo size is largely controlled by *NRPD1* or p4 (Lu et al., 2012; Kirkbride et al., 2019). This is presumably because a group of p4-siRNAs may regulate expression of their target genes in both embryo and endosperm during seed development. It will be of interest to investigate if there is a crosstalk between small RNA-mediated pathway and imprinting to regulate embryo and seed development.

## MATERIALS AND METHODS

### Plant materials and growth conditions

We utilized four lines to generate the CNS lines – Col-0, C24, ColC with the transgene *proCCA1:LUC* [Col(*CCA1*:*LUC*)], and C24C with the transgene *proCCA1:LUC* [Col(*CCA1:LUC*)] (Ng et al., 2014). The CCA1:LUC was used for an intent of diurnal studies, which were not included in this study. To generate the CNS lines, we backcrossed Col-0 with C24C over six generations, and C24 with ColC over five generations, respectively. To perform manual crosses, flowers were first emasculated by removing sepals, petals, and anthers from immature flowers, with pollination performed the following day. For example, the CNS line Col(C24C) resulted from the Col-0 in which the nuclear was replaced by C24C. We obtained Col(C24C) and C24(ColC) CNS lines. The regular reciprocal crosses were derived from Col x C24C and C24 x Col combinations. The reciprocal crosses Col x C24C and Col(C24C) x Col were used to test imprinting effect on nuclear genes in the same cytoplasmic background. At the meantime, genetic crosses Col(C24C) x Col and C24C x Col were used to test the cytoplasmic effects on gene expression and phenotypes. To investigate the role of AT4G13495, two SALK T-DNA knockout lines were obtained from the Arabidopsis Biological Resource Center – SALK 52375 and SALK 205712C.

The seeds were sterilized in 20% bleach (Clorox) for 10 minutes. Following sterilization, seeds were washed five times with 1 mL sterile ddH_2_0 and germinated on 0.5 MS media with 1% sucrose and stratified in the dark at 4°C for 48 hours. Plates were then transferred to the 22°C growth room for with 16 hours of light and 8 hours of dark per day to germinate the seeds. At 7 days after germination, seeds were transplanted onto a soil mix of three parts Pro-Mix Biofungicide to one part Field and Fairway. Soil was treated with 4g of Miracle Gro Plant Food and 1 teaspoon of Gnatrol Biological Larvicide (Valent Biosciences LLC, Mitchell County, Iowa) per gallon of water at first watering. Bonide copper soap fungicide was sprayed as needed or weekly to prevent powdery mildew infection.

### Sequencing the CNS lines

Rosette leaves were harvested from plants at 21 days after germination. Leaves were flash frozen in liquid nitrogen and stored at -80°C. Genomic DNA was extracted from the leaves using a QIAGEN Plant DNEasy mini kit (Qiagen, Hilden, Germany). DNA was sheared to a target size of ∼450-bp fragments using a sonicator (Covaris, Woburn, MA), using 4 x 14s cycles, an intensity of 4, and a 10% duty cycle, and purified using the QIAGEN PCR Purification kit. Ab aliquot of DNA (100 ng) was used to create libraries using the NEBNext Ultra II kit (New England Biolabs, Ipswich, Massachusetts). Libraries were sequenced at The University of Texas at Austin Genome Sequencing and Analysis Facility on a HiSeq 4000, with paired 150-bp reads and a total of ∼30 million reads per sample.

### Bioinformatics for genome sequences

Reads were first trimmed using fastx (http://hannonlab.cshl.edu/fastx_toolkit/) and mapped to the TAIR10 genome using bowtie (Langmead and Salzberg, 2012). The mapped reads were filtered for optical duplicates and base quality scores were recalibrated using GATK (Van der Auwera et al., 2013). Variants were called using GATK’s HaplotypeCaller and filtered to identify SNPs that had a minimum of 10 reads supporting them. A reference set of SNPs for Col-C24 SNPs was created by identifying SNPs that were called as homozygous in Col and C24, as well as heterozygous in Col x C24. Homozygous variants were filtered using GATK’s SelectVariants tool requiring that 100% of the reads supported the variant. Heterozygous variant calls were filtered using GATK’s SelectVariants to required that there was between 75%-25% allele balance between the two calls to filter out spurious heterozygous calls. Variants were filtered for depth and allele balance in the CNS lines using the same criteria, and these were then filtered against the reference set of SNPs identified in the Col, C24 and F_1_ lines. Variants per 100 kb window were counted and displayed using SnpEff (Cingolani et al., 2012).

#### Seed and embryo measurements

Plants were emasculated and then manually pollinated the following day. At 7 DAP siliques were dissected, by slicing along the replum and removing seeds with forceps (Ando et al., 2023). Seeds were placed onto a 0.5 ml drop of water, which was removed and replaced with clearing solution (chloral hydate:30% glycerol, 1:1, wt:v). Seeds were allowed to clear overnight and visualized on a Nikon Eclipse Ni compound light microscope using Nomarski optics. Seed area and embryo length were measured using ImageJ (Schindelin et al., 2012).

### Rosette size measurements

Rosettes were photographed at 21 and 28 days after germination. The rosette area and diameter were measured using PlantCV software (Gehan et al., 2017), as previously reported (Miller et al., 2015).

### Mature seed area measurements

A set of 30 seeds from each line was harvested from mature siliques and photographed using a Nikon SMZ1500 microscope. Seed area was measured using ImageJ (Schindelin et al., 2012). Seeds/embryos were manually selected along their outlines, and pixel values of selected areas were automatically generated from ImageJ after calibration from scale bars of the original pictures. Approximately 300 seeds in each cross were fixed and pictures were taken. Only seeds/embryos that were well displayed in a picture (i.e., clearly fixed and oriented to show the whole image) were chosen to be measured. A total of 20-50 images were used to estimate seed and embryo volumes (mm), respectively. Student-*t* tests were used for statistical significance analysis.

### Laser-capture microdissection (LCM)

LCM procedure followed a published protocol with two biological replicates (Ando et al., 2023). At 6 DAP, siliques were harvested by slicing along the replum and cut into 3-5 mm sections using a scalpel. The sections were immersed in a fixative of ice cold 75% ethanol and 25% glacial acetic acid. Immersed sections were put under vacuum for ten minutes to infiltrate sections with the fixative. Siliques were left immersed in the ethanol and acetic acid fixative overnight at 4°C and fixed using a microwave-based fixation method (Takahashi et al., 2010; Ando et al., 2021), in which seeds are dehydrated using an ethanol series, followed by infiltration with butanol and then paraffin. Briefly, in the following day, the fixative was replaced with fresh solution, and seeds were then microwaved at 250 W for 15 minutes in an ice bath in a PELCO Biowave 34700 with a Cold Spot (Ted Pella) set to 37°C. This was repeated three more times. Seeds were dehydrated using an ethanol series (70%, 70%, 80%, 90%, 100%, 100%, 50% butanol, 100% butanol) by replacing the existing solution with the ethanol and then microwaving the solution for 90 seconds at 350 W, with the Cold Spot set to 58°C. The seeds were microwaved in a mixture of 50% molten Paraplast Plus paraffin (Millipore Sigma, St. Louis. MO) for ten minutes at 250W. This was followed by microwaving the seeds in 100% molten paraffin, again for ten minutes at 250W. Finally, the seeds were microwaved in fresh 100% molten paraffin, partially submersed in a 75°C water bath, for 30 minutes at 250W, replacing the paraffin each time. The silique segments were then embedded into paraffin blocks, and stored at 4°C. Blocks were sectioned into 7 µm sections using a paraffin microtome and mounted onto PEN foil slides (ZEISS, Oberkochen, Germany). Slides were deparaffinized by immersion in xylenes for two minutes, followed by air-drying of the slide for five minutes. The seeds were dissected using a ZEISS PALM Laser Microdissection microscope (ZEISS, Oberkochen, Germany), isolating the embryos first, and the endosperm afterwards. Endosperm sections consisted of chalazal, micropylar and peripheral endosperm. The cut tissue was collected onto the adhesive collection cap of LCM collection tubes (ZEISS, Oberkochen, Germany). RNA was isolated from LCM-isolated tissue using the RNAqueous Micro Kit RNA isolation kit (Invitrogen, Carlsbad, California). RNA quality and quantity were checked on a Bioanalyzer 2100 using the RNA Pico eukaryotic assay (Agilent, Santa Clara, California). gDNA was removed from RNA samples using RQ1 DNAse (Promega, Madison, Wisconsin). mRNA-seq libraries were prepared using 50-100 ng of RNA (with a RIN score greater than five), using the NEB Ultra II RNA seq kit (New England Biolabs, Ipswich, Massachusetts), with the PolyA selection module. Libraries were sequenced at Genome Sequencing and Analysis Facility (GSAF) at The University of Texas at Austin, with a target of ∼20 million reads per sample. The libraries for the conventionalreciprocal crosses were sequenced using single end 100 bp reads on an Illumina HiSeq 2500. One biological replicate of the reciprocals with fixed cytoplasm was sequenced using single end 75 bp reads on an Illumina NextSeq with a target of ∼30 million reads per sample, and the second was sequenced using single end 100 bp reads on an Illumina NovaSeq with a target of ∼20 million reads per sample.

### Analysis of imprinted and differentially expressed genes

Reads were first trimmed with trimmomatic (Bolger et al., 2014) to remove low quality reads. High-quality reads were mapped onto the TAIR10 genome using STAR (Dobin et al., 2013) with the following settings: --outFilterMismatchNoverLmax 0.04 -- outFilterMultimapNmax 20 --alignIntronMin 25 --alignIntronMax 3000. Reads were filtered to identify uniquely mapped reads using samtools (-q 60 setting) (Barnett et al., 2011). For differential expression analysis, reads overlapping each gene were counted using HTseq with the union and reverse stranded settings. Gene annotations from Araport11 (Cheng et al., 2017) were used to identify gene loci. Statistical analysis of differential expression was performed using EdgeR, using linear models to identify differentially expressed genes (Robinson et al., 2010; McCarthy et al., 2012). Genes that showed a greater than two-fold change and a Bejamini-Hochberg adjusted p-value of less than 0.05 were classified as differentially expressed between the samples. To identify imprinted genes, bam files of uniquely mapped reads were processed using a previously published python script that discriminated variants using bam files. The read counts from the called variants and a bed file indicating the location of known SNPs that overlap with genes were used to estimate the number of reads corresponding to the paternal and maternal origins at each gene (Wyder et al., 2019). The EdgeR protocol utilizing linear models to identify imprinted genes was used for statistical analysis. Genes with a Benjamini-Hochberg adjusted p-value of less than 0.05 were identified as imprinted genes for further analysis.

GO enrichment analysis was performed using the TopGO package (https://bioconductor.org/packages/release/bioc/html/topGO.html) using the weight01 algorithm. Significantly enriched GO terms were defined as having a p-value lower than 0.05.

### Analysis of cross-specific imprinted genes

Cross-specific imprinting of genes was identified using the workflow outlined in a published paper (Pignatta et al., 2014). Briefly, log_2_(maternal/paternal) read ratios were calculated and plotted for all pairs of reciprocal crosses. These were then filtered for genes that fell within a Euclidian distance of 1 from x = 1 or y = 1, which identifies genes that lack parental bias in at least one parent. A normalized parental bias factor was calculated for all remaining genes, and genes within the top 5% of parental bias were retained. Genes that had the same parental bias in both replicates, and fell within the top 5% parental bias, were classified as having cross-specific expression in one reciprocal cross, but not in the other. These genes were compared with the imprinted genes identified previously (Gehring et al., 2011; Hsieh et al., 2011; Pignatta et al., 2014) to identify cross-specific imprinted genes, which display imprinting in crosses with other ecotypes, but lacked imprinting in only one of the reciprocal crosses (Col X C24 or C24 X Col).

### RT-qPCR validation of gene expression

RNA was extracted from three pools of ten seedlings at 7 days after germination per sample. The seedlings were flash frozen in liquid nitrogen, and the ground to a fine powder in a QIAGEN TissueLyser LT (Qiagen, Hilden, Germany). RNA was extracted from the powder using TRIzol reagent (Invitrogen, Carlsbad, California), and resuspended in DEPC-treated ddH_2_O. An aliquot of RNA (1 µg) was treated with RQ1 DNAse (Promega, Madison, Wisconsin), and 500 ng of RNA was then reverse transcribed using the Omniscript RT kit (Qiagen, Hilden, Germany). The cDNA from this reaction was used as template for qPCR, using the FastStart Universal SYBR Green Master Mix (Roche, Indianapolis, Indiana). The following PCR cycle was used for all RT-qPCR experiments: 50°C, 120s preincubation and a 50°C, 600s secondary preincubation followed by the following two-step amplification cycle: 40 cycles of 95°C for 15s, followed by 60°C for 60s. The LightCycler 96 system (Roche, Indianapolis, Indiana) was used for qPCR, and relative expression levels of each gene were calculated using the LightCycler software and *ACT7* (AT5G09810) as a control. Controls without reverse transcriptase were performed for each sample, and two technical replicates were performed for each of the three biological replicates per sample. Primers used for RT-qPCR can be found in Supplementary Table 1.

### Stem-loop RT-qPCR for analysis of miRNA relative expression

Stem-loop reverse transcription and qPCR was adopted from a published protocol (Varkonyi-Gasic, 2017). The U6 snRNA used as a control for relative expression of miR863. Briefly, RNA was extracted from three pools of ten seedlings at 7 days after germination per sample. The seedlings were flash frozen in liquid nitrogen, and the ground to a fine powder in a QIAGEN TissueLyser LT (Qiagen, Hilden, Germany). RNA was extracted from the powder using TRIzol reagent (Invitrogen, Carlsbad, California), and resuspended in DEPC-treated ddH_2_O. 50 ng of RNA was used in a pulsed reverse transcription reaction performed using SuperScript III RT (Invitrogen, Carlsbad, California), with separate reactions for U6 snRNA and miR863. Reverse transcription was carried out at 16°C for 30 min, followed by 60 cycles of 30°C for 30s, 42°C for 30s and 50°C for 1s. The reaction was then inactivated by incubation at 85°C for 5 min. cDNA was then used as a template for qPCR using the Luna Universal Probe Master Mix (New England Biolabs, Ipswich, Massachusetts) with the Universal Probe Library #21 probe (Roche, Indianapolis, Indiana). qPCR was performed with one 95°C incubation for 5 minutes, followed by 50 two-step cycles of 95°C for 5s and 60°C for 10s. The LightCycler 96 system (Roche, Indianapolis, Indiana) and its associated software were used to perform qPCR and quantification of the results. Controls without reverse transcriptase were performed for each sample, and two technical replicates were performed for each of the three biological replicates per sample. Primers used for RT-qPCR can be found in Supplementary Table 1.

### Knockout of *HDG9* using CRISPR-Cas9

Two gRNAs were designed to target HDG9 (AT5G17320) using the CRISPR-PLANT tool (https://www.genome.arizona.edu/crispr/instruction.html) and the CRISPR-P v. 2.0 tool (http://crispr.hzau.edu.cn/CRISPR2/). A construct containing Cas9 and a cassette of the two gRNAs was assembled using the Golden Gate assembly protocol outlined in (Xing et al., 2014), using the pHEE401E construct from (Wang et al., 2015). The constructs were used to transform *A. tumefaciens* GV3101, which was used to transform the Col and C24 accessions, respectively, by floral dip (Clough and Bent, 1998).

## Data availability

RNA sequencing data were deposited under the Gene Expression Omnibus accession no. GSExxxx.

## Acknowledgements

We thank Dr. Alan Lloyd at The University of Texas at Austin for supervision in the latter part of this project and Texas Advanced Computing Center for providing computing support for data analysis. The financial support for this work was partly provided by the National Institutes of Health (GM109076) to ZJC.

## Author contributions

V.J. and Z.J.C. conceived the research, analyzed the data, and wrote the paper. X.S. performed the experiments.

## SUPPLEMENTAL FIGURES

**Figure S1.** Genomic analysis of CNS lines showing nuclear genome conversion while retaining their original mitochondrial and plastid genomes.

**Figure S2.** Seedling growth vigor observed between CNS reciprocal hybrids with fixed cytoplasmic genomes, but not in CNS reciprocal hybrids with fixed imprinting.

**Figure S3.** Tissue contamination was absent in the samples collected via LCM.

**Figure S4.** Weak expression of the imprinted genes only in the conventional reciprocal crosses or in CNS reciprocal hybrids with fixed cytoplasm.

**Figure S5.** Analysis of cross-specific imprinted genes using LCM samples.

**Figure S6.** Seed phenotypes of CRISPR/Cas9-edited *HDG9* lines.

**Figure S7.** A substantial more differentially expressed genes in the CNS reciprocal hybrids with fixed cytoplasm than those in the CNS reciprocal hybrids with fixed imprinting.

**Figure S8.** Endosperm-specific expression of all embryonic imprinted genes except *AT4G13495*.

**Figure S9.** Analysis of imprinted AT4G13495 locus consisting of two miRNAs in T-DNA insertion lines.

**Figure S10.** Embryo and seed size is affected by *NRPD1*.

## SUPPLEMENTARY DATA

**S1. Primers used for PCR amplification**

**S2. Imprinted genes in the endosperm**

**S3. Genes with cross-specific imprinting**

**S4. Differentially expressed genes between reciprocal hybrids in the endosperm**

**S5. Differentially expressed genes between reciprocal hybrids in the embryo**

**S6. Imprinted genes in the embryo**

